# Maf is a regulator of differentiation for gut immune epithelial cell Microfold cell (M cell)

**DOI:** 10.1101/2021.10.15.464565

**Authors:** Joel Johnson George, Fábio Tadeu Arrojo Martins, Laura Martin-Diaz, Keijo Viiri

**Affiliations:** Faculty of Medicine and Health Technology, Tampere University Hospital, Tampere University Tampere, Finland

**Keywords:** Organoids, *in vitro* modeling, Gut immunity, M cells, Peyer’s patch, Maf

## Abstract

Microfold cells (M cells) are a specialized subset of epithelial intestinal cells responsible for immunosurveillance of the gastrointestinal tract. M cells are located in the Peyer’s patches and are crucial for monitoring and the transcytosis of antigens, microorganisms, and pathogens via their mature receptor GP2. A mature M cell with Gp2 receptor aids in the uptake of antigens, which are passed through the single layer of epithelium and presented to underlying antigen-presenting cells and processed further down-stream with B cells, T cells, and dendritic cells. Recent studies revealed several transcription factors and ligands responsible for the development and differentiation of mature M cells however, an exhaustive list of factors remains to be elucidated. Our recent work on the epigenetic regulation of M cell development found 12 critical transcription factors that were controlled by the polycomb recessive complex 2. Musculoaponeurotic fibrosarcoma transcription factor (Maf) was identified as a gene regulated by the polycomb repressive complex (PRC2) during the development of M cells. In this paper, we explore Maf’s critical role in M cell differentiation and maturation. Maf falls under the purview of RANKL signaling, is localized in the Peyer’s patches of the intestine, and is expressed by M cells. Given that, complete knockout of the Maf gene leads to a lethal phenotype, organoids isolated from Maf knockout mice and treated with RANKL exhibited impaired M cell development and a significant decrease in Gp2 expression. These findings reveal that Maf is an important regulator for M cell development and differentiation.

## 1. Introduction

The mucosal lining of the gastrointestinal tract is a constant battlefront where intestinal tissues are up against microbes, viruses, antigens, and other harmful pathogens. One of the defense mechanisms it employs is the immune-inductive sites that are found in the gut-associated lymphoid tissue also known as Peyer’s patches (PP). Peyer’s patches compose of three distinct regions; a germinal center (GC), an interfollicular region abundant in T cells and the sub-epithelial dome (SED) which houses lymphoid cells such as B cells, T cells, and other antigen-presenting cells ^1–3^. Since the PP’s lack a lymphatic system where antigens could potentially be transported, the PP’s directly sample mucosal antigens through specialized immune epithelial cells, these cells are known as Microfold Cell (M cell) ^2,4^. M cells are phagocytic epithelial cells that enable the uptake and transcytosis of luminal antigens into the gut-associated lymphoid tissue (GALT), they are responsible for the rapid transport of bacterial antigens to antigen-presenting immature dendritic cells ^5–7^. Transcytosis is achieved through their surface receptor Glycoprotein 2 (GP2) which binds to antigens and aids in uptake, a mature M cell is characterized by the presence of a functioning Gp2 ^8,9^. M cells form a part of the adaptive immune system as they provide residence to B cells, T cells, and dendritic cells. They are uniquely different in morphology compared to neighboring epithelial cells as they are characterized by short and irregular microvilli and a lack of mucus layer to aid in transcytosis. they have pocket-shaped invagination under which B cells, T cells, macrophages, and dendritic cells are present ^10–13^.

M cell differentiation is dependent on the stimulation of Receptor activator of nuclear factor κB ligand (RANKL) which is a member of the tumor necrosis factor (TNF) family cytokine ^14,15^. Cycling intestinal stem cells in the crypts that express Lgr5 and Rank receptors undergo differentiation to M cells after exposure to RANKL which is secreted by stromal cells under the follicle-associated epithelium (FAE) known as M cell inducer cells (MCi) ^16^. Recent studies demonstrated that Spi-B and Sox8 are transcription factors necessary for M cell development and differentiation. Spi-B null mice lacked transcytosis capacity due to the lack of M cells and demonstrated an impaired mucosal response to *S. Typhimurium* ^15,17^. Sox8 null mice gave rise to an immature phenotype of M cells without Gp2 receptor thereby indicating a loss in immune response in the PP’s of mice orally infected with *S. Typhimurium*^18^. However, Sox8 expression is active in Spi-B null mice and Spi-B is actively transcribed in Sox8 KO mouse yet both sets of mice show a lack of Gp2 receptor, impaired mucosal response and a severe lack of transcytosis capacity ^18^. This implies that there are additional transcription factors involved in M cell development and they remain to be characterized.

Recently, we published our work looking at the epigenetic regulation of M cell differentiation and found that the polycomb repressive complex 2 (PRC2) regulates 12 transcription factors critical for M cell development. PRC2-regulated Estrogen related receptor gamma (Esrrg) was necessary for Sox8 expression and Gp2 expression for mature M cells ^19^. Atonal BHLH Transcription Factor 8 (Atoh8), another transcription factor regulated by PRC2, negatively regulated the M cell population in the Peyer’s patches; Atoh8 null mice showed an increase in M cell population and a higher transcytosis capacity ^20^. Maf was another gene identified to be regulated by the PRC2 complex. Maf has been shown to be critical for the development and differentiation of various signaling mechanisms with tissues such as the cornea and lens development, mechanoreceptors involved in touch sensations, differentiation of chondrocytes ^21–24^. Maf has been identified as an immune regulator and a transcription factor for T helper 2 cells (Th2), additionally, it has also been found to play a key role in the differentiation and development of innate immune cell types, B lymphocytes, and T-cell subsets in Peyer’s patches ^25–30^. Our findings here identify Maf to be an additional regulator of M cell differentiation that is dependent on Rank-RANKL signaling. Maf expression was localized to the Peyer’s patch and necessary for the expression of early markers of M cell differentiation and expression of mature M cell marker Gp2. These observations indicate that Maf is a novel player essential for the differentiation and development of M cells in the follicle-associated epithelium.

## 2. Results

### 2.1. Maf is regulated by the PRC2 and localized in the Peyer’s patch

Our recent data from the chromatin immunoprecipitation assays with sequencing (ChIP-seq) and global run-on sequencing (GRO-seq) analysis of M cells (RANKL treated cells) and crypt/ISC (WENRC treated cells) revealed how the epigenome regulates the differentiation and development M cells in the gut^19^. Furthermore, PRC2-regulated *Maf* turned up as one of the genes highly expressed in the analysis (log2 fold change -2.66 RANKL vs WENRC (crypt/ISCs) (Fig.1A). Leveraging the Maf LacZ mouse model where the Maf gene is disrupted by positioning LacZ gene in the Maf locus, we traced the gene expression of Maf in the GALT of Maf heterozygous mice through β-galactosidase activity^31^. The staining revealed Maf expression to be localized in the Peyer’s patch (Fig. 1B). Organoids treated with RANKL underwent immunofluorescence staining of GP2 and Maf. The confocal images revealed their expression to be localized together (Fig. 1C).

**Figure 1.**
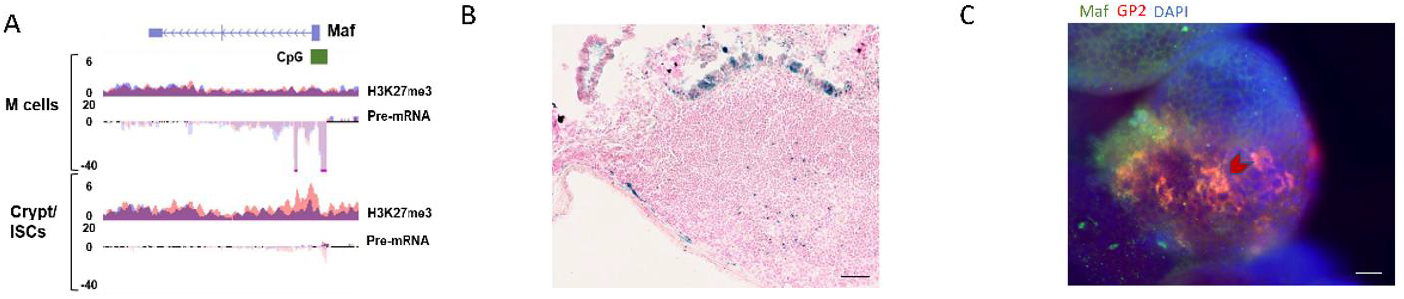
Maf is PRC2 regulated and localized in the FAE. A) H3K27me3 occupancy at CpG islands spanning the promoter and first exon of the Maf gene in organoids treated with RANKL (M cells) and treated with Wnt3a and Chir99021 (Crypt/ISCs). Below, pre-mRNA expression of Maf in organoids treated as above (y-axis: normalized tag count, ENR500 = R-spondin 500ng/ml, R100 = RANKL 100ng/ml). Original data published in George et al. 2021 CMGH 10.1016/j.jcmgh.2021.05.014 B) Sections of Peyer’s Patch from Maf heterozygous mice were processed and stained with β-galactosidase staining assay to trace Maf expression in the FAE. Bar, 100 µm C) immunofluorescence of GP2 and Maf in RANKL treated organoids, Bar, 100 µm.

### 2.2. Maf expression is induced by RANKL stimulation and is under the Rank-RANKL signaling axis

To investigate if Maf is expressed by RANKL stimulation, intestinal organoids derived from wildtype mice were cultured with and without RANKL treatment for 3 days. qPCR analysis of isolated RNA shows significant upregulation of *Maf* as well as *Gp2* in organoids treated with RANKL. (Fig.2A) β-galactosidase experiments to trace Maf expression in Maf heterozygous organoids grown in RANKL conditions revealed Maf expression in cells leading to the lumen of the organoids (Fig.2B). Rank-KO mouse intestinal organoids were generated using the Lenti V2 CRISPR/Cas9 system, RANKL treatment of RANK KO did not activate *Maf* expression suggesting that Maf falls under the control of Rank-RANKL signaling (Fig. 2C).

**Figure 2.**
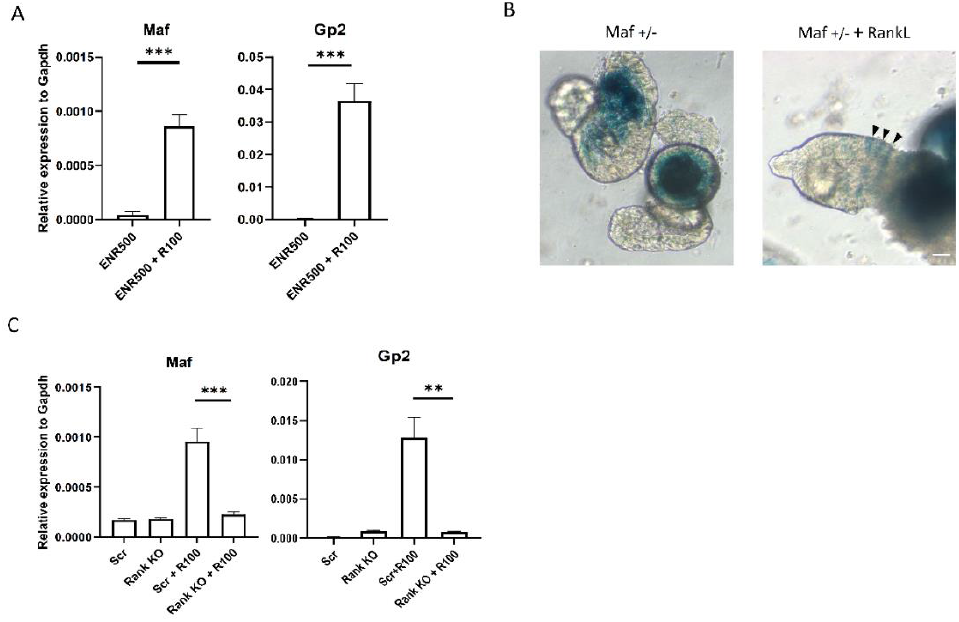
Maf expression in M cells is dependent on Rank- RANKL signaling A) Organoids generated from wild-type mice were stimulated with 100ng of RANKL for 4 d. B) Organoids isolated from Maf heterozygous mice were processed into paraffin blocks and stained with β-galactosidase staining assay to trace Maf expression in the organoids. Arrowheads indicate Maf expressing M cells. Bar, 100 µm C) Rank KO organoids and Scrambled organoids generated by lentiCRISPR v2 were incubated with RANKL for 4 days, *Maf* and *GP2* expression was analyzed by RT-qPCR. In (A&C) unpaired two-tailed Student’s t-test was performed for three independent experiments, ^***^, P < 0.005; ^**^, P < 0.01.

### 2.3. Maf deficiency impairs M cell differentiation and other M cell-associated factors

Given that Maf is prominently expressed in Peyer’s patches, we looked to see if the abolition of Maf had any effect on the development of M cells. Complete knockout of the Maf gene produce a lethal phenotype and pups were born still-born or only survived for up to 4 hours. However, organoids were isolated from both Maf WT and Maf KO pups as soon as the mothers went into labor and cultured in media conditioned with Egf, Noggin, and R-spondin, Wnt, and Chir99021. After propagation, the organoids were grown in the presence and absence of RANKL for 3 days. Rt-qPCR analysis of the isolated RNA reveals a significant decrease in *Gp2* expression; *SpiB, Sox8*, and *Esrrg*, transcription factors that are critical for functionally mature M cells showed significantly reduced expression as well (Fig.3A). Early developmental markers of M cell differentiation such as *MarcksL1* and *Tnfaip2* showed diminished expression in the Maf knockout organoids compared to its wildtype counterparts. Immunofluorescence analysis of Gp2 in Maf WT and Maf KO depicted a lack of Gp2 expression. Our observations reveal that Maf is required for the expression of early markers as well as maturation genes associated with M cell differentiation.

**Figure 3.**
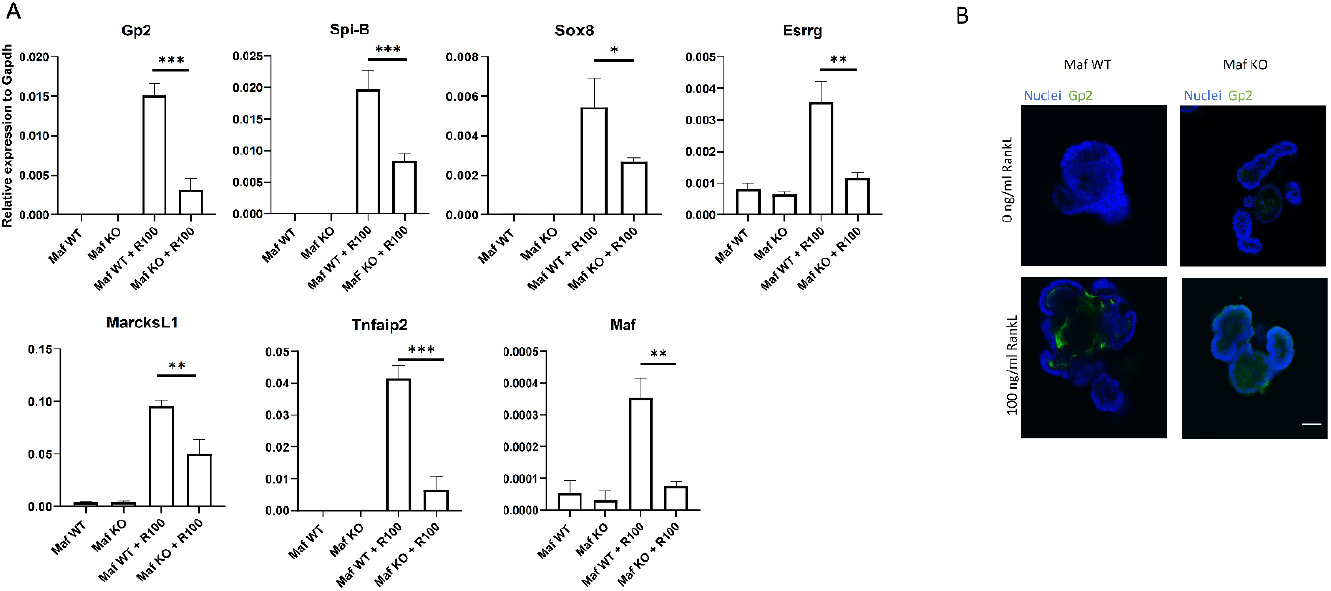
Impaired M cell maturation in Maf KO results from epithelium intrinsic defect. A) qPCR analysis of M cell-associated genes from RNA of organoids derived from Maf WT and Maf KO mice cultured with and without RANKL for 3 d. Values are presented as the mean ± SD; ^***^, P < 0.005; ^**^, P < 0.01; ^*^, P < 0.05; unpaired two-tailed Student’s t-test, n = 3. Data are representative of two independent experiments. (B) Immunostaining images for GP2 (green) on organoids. Bar, 100 µm.

## 3. Discussion

M cells are specialized intestinal epithelial cells that are a part of the adaptive immunity and on the frontlines of the gut epithelium initiating a mucosal immune response against pathogenic bacteria. These epithelial cells partake in gut immunity by transcytosis of pathogens and commensal bacteria via the receptor Gp2. Recent research into M cells has revealed critical transcription factors such as Spi-B, Sox8, Esrrg, and Atoh8 and various other ligands like membranebound RANKL from lymphoid cells and S100A4 from Dock8 cells essential for M cell development. ^14,17–19,32–34^ However, complete elucidation of the differentiation and development of M cells remains to be understood. Recently we explored the epigenetic regulation of M cells and the analysis revealed several PRC2-regulated transcription factors controlling M cell development and differentiation (6 silenced and 6 upregulated). Maf was found upregulated in the list of genes identified^19^. From this study, we describe that M cells in the FAE express Maf and are under the overview of RANKL signaling and critical for the maturation of functional M cells.

Maf (musculoaponeurotic fibrosarcoma) gene encodes for the transcription factor Maf (c-Maf). The Maf transcription family comprises of 7 members divided into 2 subclasses based on their size; large Maf proteins include MAFA/L-MAF, MAFB, MAF/c-Maf, and NLR (neural retina leucine zipper), and the small Maf proteins, MAFK, MAFG, and MAFF. The small Maf proteins are also distinct by the lack of amino-terminal transactivation domain. The Maf family is highly conserved and has a unique leucine zipper structure (bZIP). In the pancreas, c-Maf regulates glucagon hormone production and in the liver, it is critical for erythropoiesis^35–37^. Maf also plays a role as an immune regulator and a transcription factor necessary for T helper 2 cells (Th2), additionally, it has also been found to play a key role in the differentiation and development of innate immune cell types, B lymphocytes, and T-cell subsets ^25–28^. In line with Maf being a major contributor to cellular, signaling, and physiological process, Maf knockout mice with C57BL/6 background are lethal embryonically or perinatally ^21,22,36^. They are often born stillborn or only live up to 4 hours. Maf KO with BALB/c background lives to adulthood.

Our work here characterizes the role of Maf in M cell differentiation and development. X-gal assay of Maf-LacZ heterozygous mice show Maf expression in the Peyer’s patch of the follicle-associated epithelium and organoids isolated from the Maf heterozygous mice showed that Maf was expression was inducible upon RANKL stimulation. Maf tracing via X-gal assay showed Maf expressing cells away from the crypt and closer towards the lumen in organoids treated with RANKL. As expected, pups with complete knockdown of Maf were stillborn or only lived 2-4 hours after birth but intestinal crypts isolated and cultured in RANKL were able to provide insights into Maf’s critical role in M cells development. Maf deficit organoids showed a significant decrease in the maturation of M cells as Gp2 expression was significantly reduced indicating impaired and immature M cells. Lack of Gp2 expression in M cells have been shown to result in attenuation of antigen sampling and transcytosis and increased rates of infection to *S. typhimurium*. Since this conclusion of impaired M cell development was reached using epithelial organoids, it can also be assumed that loss of Gp2 in Maf KO organoids is an epithelium-intrinsic defect. Key transcription factors required for M cell maturation such a*s Spi-B, Sox8*, and *Esrrg* showed absent or reduced activation without Maf. Early developmental markers of M cells such as *MarcksL1* and *Tnfaip2* were also affected and showed reduced activation. In line with key transcription factors and early developmental factors showing absent or reduced expression, it is reasonable to conclude that Maf is a critical factor for M cell differentiation. In ocular lens differentiation and osteogenic differentiation Maf expression is regulated by BMP signaling ^38,39^. Atoh8, a transcription factor critical for maintaining the density of the M cell population was also found to be under the regulation of BMP signaling ^32^. It is possible that RANKL induced BMP signaling could be responsible for regulating the expression of Maf in M cells. However, further experiments with Maf KO in mice with BALB/c background are required to understand the physiological significance of Maf KO in-vivo.

In conclusion, PRC2 regulated Maf has major implications in M cell maturation and mucosal immunity. The transcription factor is necessary for M cell development and lack of Maf could lead to attenuation of transcytosis of antigens and commensal bacteria. Reduced transcytosis has been shown to lead to an impaired immune response to pilli-expressing pathogenic bacteria such as *E*.*coli* and *S. typhimurium*^40^.

## 4. Materials and Methods

### 4.1. Mice

All animal experiments were approved by the Finnish National Animal Experiment Board (Permit: ESAVI/5824/2018). Maf lacZ mice were caged in standard light-dark conditions at the pathogen-free animal facility of the faculty of Medicine and Health Technology. Food, water ad libitum was followed in a regularly timed schedule. B6.129-Maftm1Gsb/J heterozygous mice were purchased from Jackson laboratories (Cat number: 004158 | Maf^lacZ^). To generate Maf wildtype line, heterozygous line, and homozygous line, B6.129-Maftm1Gsb/J heterozygous were mated with, B6.129-Maftm1Gsb/J heterozygous, F1 generation was backcrossed with heterozygous mice. Littermates with Maf wildtype were used as control. Maf genotypes were confirmed by qPCR.

### 4.2. β-. galactosidase staining of Peyer’s patches and organoids

Ileal PP’s were isolated from the ileal section of the small intestine and transferred to a 10cm dish with 30ml of cold PBS. Excess fat was cut from the tissues and embedded into paraffin blocks. Organoids grown in RANKL was washed with PBS and made into paraffin blocks. The blocks were cut at 10um sections and mounted on a slide. X-gal (5-Bromo-4-chloro-3-indoxyl-beta-D-galactopyranoside, Goldbio) was dissolved in dimethylformamide at 50 mg/ml. Paraffin blocks were fixed with 4% PFA for 10 minutes. The slides were washed 3 times with 3 changes of PBS for 5 minutes wash and the final rinse in distilled water. After drying the slides were incubated in X-gal working solution at 37 C for 24 hours in a chamber with adequate moisture content. The sections were washed in PBS solution 2 times for 5 minutes each. After rinsing with distilled water, the sections were counter-stained with nuclear fast red for 3-5 minutes followed by further rinsing and washing in distilled water for 2 minutes. The sections were finally dehydrated for 3 minutes each in 70%, 95%, and twice with 100% ethanol and thrice with xylene. The sections were mounted Permount and covered with a coverslip and examined by digital slide scanner Hamamatsu Nanozoomer.

### 4.3. Mouse Intestinal Organoid culture

Mouse intestinal crypts were isolated and cultured in an *in vitro* setting as previously described ^15,41^Crypts were isolated as soon as the stillborn pups were delivered. Isolated duodenums were washed in PBS and longitudinally cut. Villi were gently scraped with a glass slide. Following washes with PBS, the tissue was cut into 2mm pieces and pipetted up and down with a 10 ml pipette. After repeated changes of the PBS till the suspension was clear, the tissues were suspended in 10mM EDTA in PBS for 20 minutes rocking at room temperature. The crypts were separated from the rest of the tissues using a 70-μm cell strainer (Fisher Scientific). The crypts were counted and cultured on a 24 well plate by embedding them in 30ul of cold Matrigel (Corning). Organoids were cultured in an optimal medium consisting of advanced DMEM/F12 (Thermo Fisher Scientific) that contained HEPES (10mM, Sigma-Aldrich), Glutamax (2mM, Thermo Fisher Scientific), Penicillin-streptomycin (100U/ml, Sigma-Aldrich), B-27 supplement minus Vitamin A (Thermo Fisher Scientific), N-2 supplement (Thermo Fisher Scientific), N-acetylcysteine (1 mM; Sigma-Aldrich), recombinant murine EGF (50 ng/ml; Thermo Fisher Scientific), recombinant murine Noggin (100 ng/mL; PeproTech), recombinant mouse R-spondin1 (1 μg/mL; R&D Systems). Media were changed every 2 days. For M cell differentiation, recombinant mouse RANKL (100ng/ml, Peprotech) was added to the media and incubated for 4 days.

### 4.4. CRISPR–Cas9 gene knockout of intestinal organoids

Guide RNAs for gene encoding RANK were designed using CRISPR design tool (http://crispr.mit.edu)^42^. These were cloned into the vector lentiCRISPR v2 (Addgene, 52961). The cloned product was transfected into HEK 293FT cells (ThermoFisher R7007). The supernatant was collected after 48 h and the Lenti-X concentrator (Clontech) was added to the suspension. The 293FT cell line was tested for mycoplasma. Intestinal organoids were cultured in ENCY (EGF, Noggin, Chir-99021, and Y-27632) prior to transduction. After the organoids were dissociated into single cells using TrypLE Express (Thermo Fisher Scientific) and supplemented with 1,000 U/ml DnaseI at 32 °C for 5 mins, the cells were washed once with Advanced DMEM and resuspended in a transduction medium (ENR media supplemented with 1mM nicotinamide, Y-27632, Chir99021, 8 μg/ml polybrene (Sigma-Aldrich)) and mixed with the previously collected concentrated virus. The mixture was centrifuged at 600 x g 32 °C for 1 hr followed by 3 hr incubation at 37°C, after which they were collected and plated on 60% Matrigel with enriched transduction medium without polybrene. On day 2 and day 4, RANK transduced organoids were selected with 2 μg/ml of puromycin (Sigma-Aldrich). Surviving clones were expanded in cultured with ENR medium. RANK KO organoids were confirmed by western blot to validate the expression of deleted gene.

### 4.5. Immunofluorescence of Organoids

Maf WT and Maf KO Intestinal crypt organoids were analyzed by whole-mount immunostaining. The organoids were cultured for 4 hours in an 8-well chamber plate in the presence and absence of RANKL 100ng/ml following fixation with 4% PFA for 15 minutes, followed by permeabilization with 0.1% Triton X-100 for another 15 minutes. The organoids were stained with Gp2 (MBL, D278-3) antibodies overnight at +4 degrees Celsius. This was followed by 2-hour incubation of anti-Rabbit secondary for Gp2 (and Anti-Rat for the secondary antibody. Gp2 expressing cells were analyzed by Nikon A1R+ Laser Scanning Confocal Microscope after mounting with ProLong Diamond with Dapi mounting solution (Molecular Probes P36962).

## Author Contributions

JJG, KV: Study concept and design. FTAM: Generation of Maf WT, heterozygous and homozygous mice and crypt isolation. JJG, LMD: staining, JJG RNA analysis and qPCR analysis. JJG: experiments and acquisition of data. JJG, KV: analysis and interpretation of data. JJG, KV: manuscript drafting. KV: project administration and funding acquisition. JJG, LMD, KV: critical revision of the manuscript for important intellectual content. All authors approved the final version of the manuscript.

## Abbreviations

M cells: Microfold cells
PP: Peyer’s Patch
FAE: Follicle associated epithelium
VE: villous epithelium
RANKL: Receptor activator of nuclear factor kappa B ligand
Rank: Receptor activator of nuclear factor kappa B
PRC2: polycomb repressive complex 2
Maf: Musculoaponeurotic fibrosarcoma

---

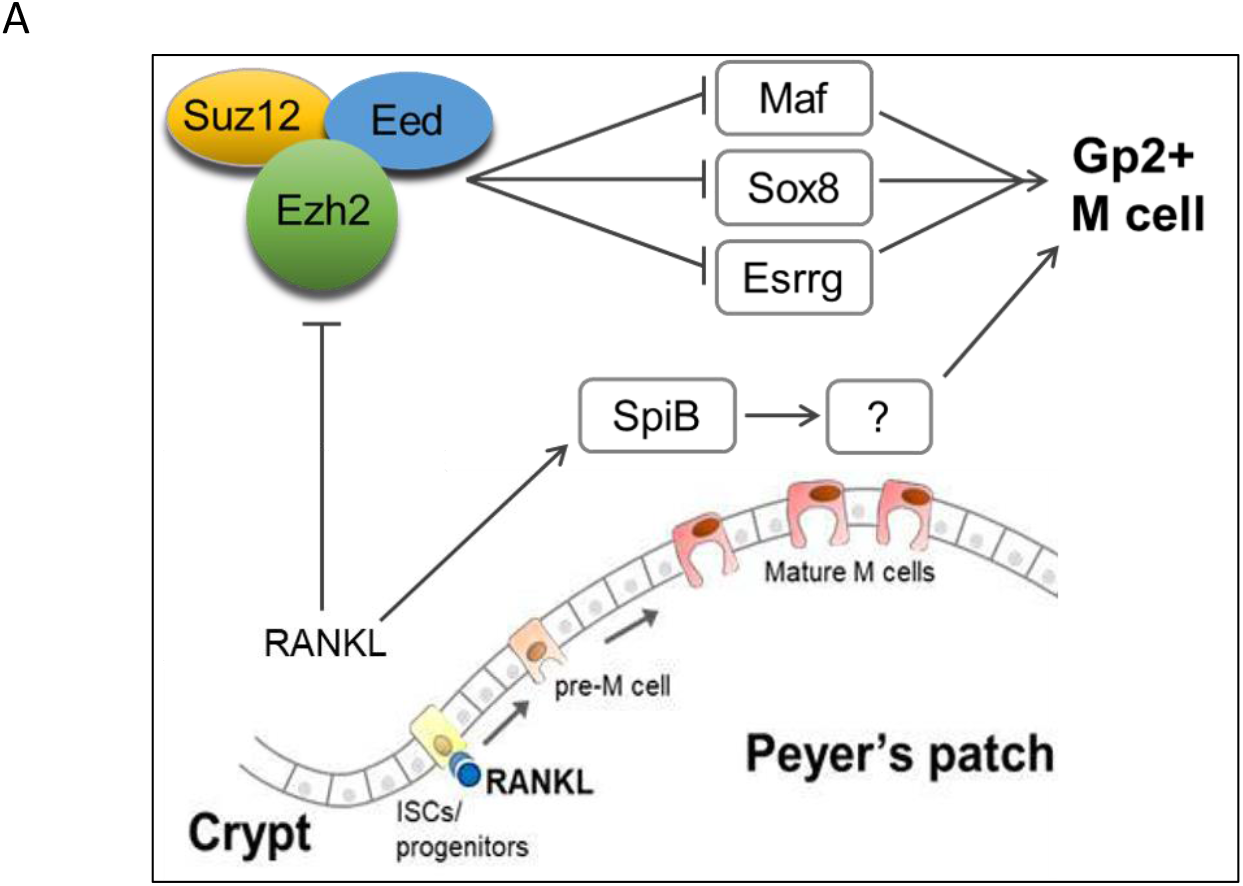

